## supplementary information for "Plasticity of growth laws tunes resource allocation strategies in bacteria"

Mukherjee et al.

### Supplementary Information

#### Supplementary Figures

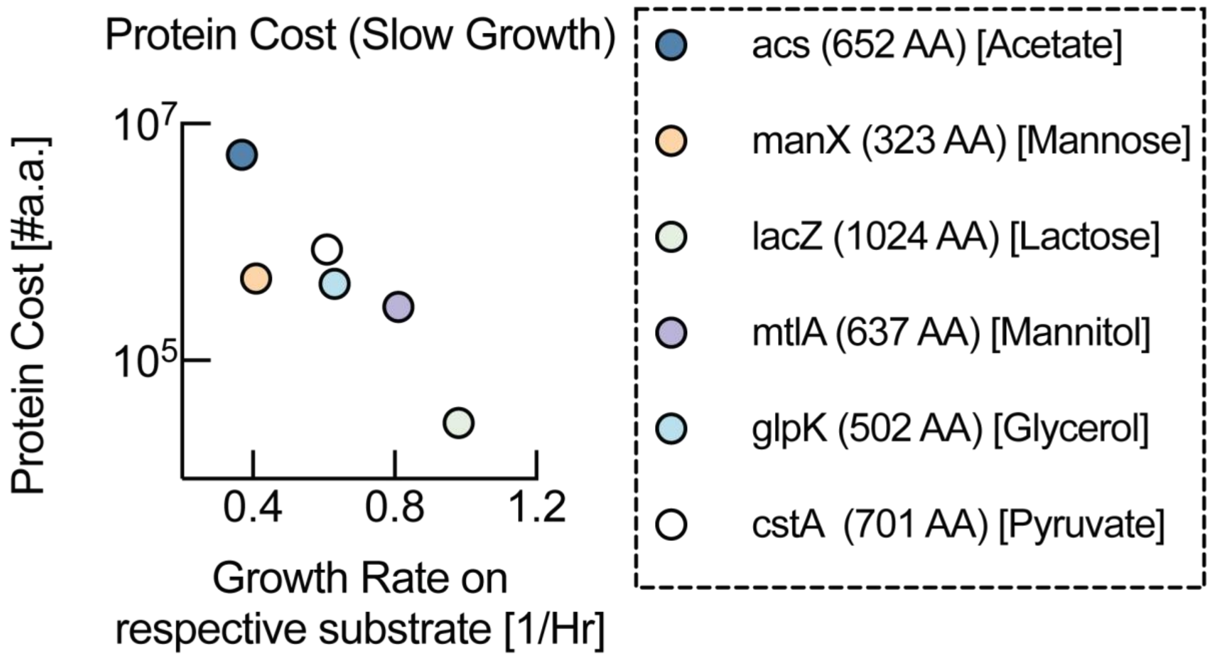

***Fig. S1: Protein cost of substrate-specific transporters and enzymes.*** *Protein copy number can be converted to protein cost in units of number of amino acids by multiplying copy number with the number of amino acids in each protein. A similar inverse correlation as the one observed for copy number in Fig. 1b of the main text, also holds for protein cost.*

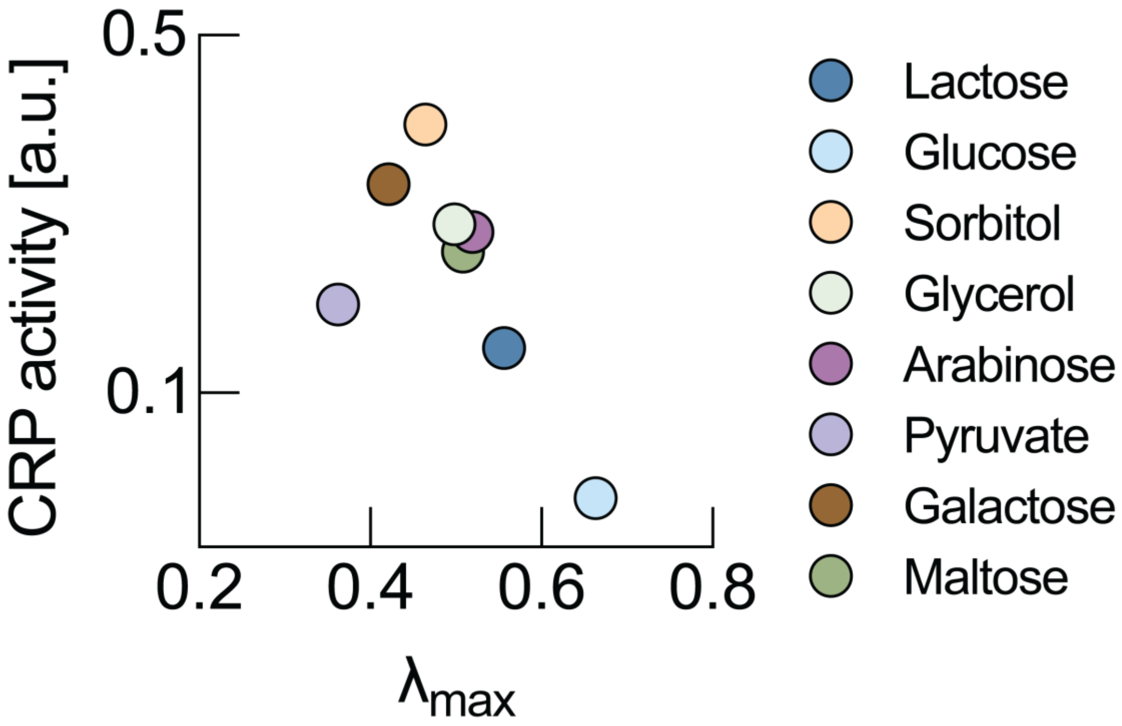

***Fig. S2: Crp activity at maxium growth rates for cAMP titrations.*** *In Fig. 2e of the main text, we plotted endogenous CRP activity measured by Towbin et al.* [1] *against growth rate on the respective substrate. Endogenous CRP activity and CRP activity where growth rate is maximum are not always the same as shown by Towbin et al.*[1]*. Therefore, we plot CRP activity at maximum growth rate against maximum growth rate from Towbin et al.*[1] *here and find a similar relationship.*

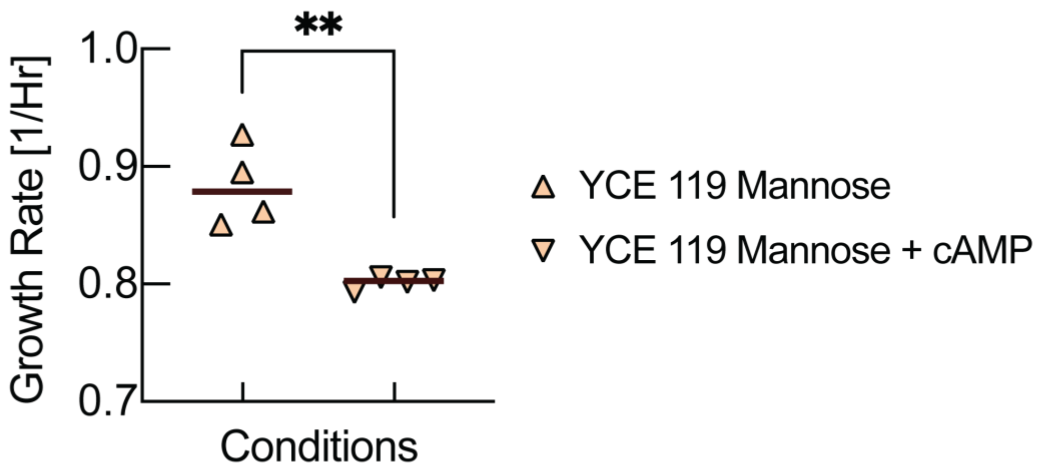

***Fig. S3: Growth rate of swapped promoter strain with the addition of cAMP.*** *As shown in the main text, many adaptability phenotypes that are lost in the swapped promoter strain (YCE 119) can be rescued by the addition of cAMP (3.5mM) to the growth medium, which upregulates the AD-sector. However, these improvements come at the cost of a reduced growth rate due the substantial protein cost of the AD-sector.*

### Supplementary Tables

| **Table S1:** | | | | | |
|---|---|---|---|---|---|
| | **Copy numbers in MOPS minimal media (from Li et. al.)** | **Number of amino acids** | **Calculated fold change at 0.45 growth rate (from Hui et. al.)** | **Copy number (at growth rate 0.45)** | **Protein cost (at growth rate 0.45)** |
| **Calculation key→** | **X** | **Y** | **Z** | **X • Z** | **(X • Z) • Y** |
| **Gene** | | | | | |
| **acs** | 643 | 652 | 13.10 | **8423.06** | **5491832.64** |
| **manX** | 2117 | 323 | 0.72 | **1527.21** | **493287.88** |
| **lacZ** | 10 | 1024 | 2.90 | **28.98** | **29679.64** |
| **mtlA** | 197 | 637 | 2.25 | **442.90** | **282129.51** |
| **glpK** | 129 | 502 | 6.84 | **881.85** | **442688.25** |
| **cstA** | 177 | 701 | 6.97 | **1233.16** | **864444.46** |

***Table S1: Protein copy number calculation.*** *Protein copy number calculation of the main carbon transporting enzyme (or the first enzyme in the primary carbon degradation pathway) from Li at. al.* [2].

| Table S2 | | | |
|---|---|---|---|
| | **Fold change calculation from Hui et. Al.** | | |
| | **0.45** | **1.04** | **←Growth rate** |
| | **a** | **e** | **←Expression level** |
| **Calculation key→** | **a/e** | **e/e** | **←Fold change** |
| **Gene** | | | |
| | 9.98 | 0.76 | ←expression level |
| **acs** | **13.10** | 1 | ←Calculated fold change |
| | 0.78 | 1.08 | ←expression level |
| **manX** | **0.72** | 1 | ←Calculated fold change |
| | 2.44 | 0.84 | ←expression level |
| **lacZ** | **2.90** | 1 | ←Calculated fold change |
| | 2.22 | 0.99 | ←expression level |
| **mtlA** | **2.25** | 1 | ←Calculated fold change |
| | 4.95 | 0.72 | ←expression level |
| **glpK** | **6.84** | 1 | ←Calculated fold change |
| | 6.23 | 0.89 | ←expression level |
| **cstA** | **6.97** | 1 | ←Calculated fold change |

***Table S2: Protein copy number fold changes across growth rates.*** *Fold change calculation of the main carbon transporting enzyme (or the first enzyme in the primary carbon degradation pathway) at slow growth rate (0.45) from Hui. et. al.* [3].

### Supplementary References

1. Towbin BD, Korem Y, Bren A, Doron S, Sorek R, Alon U. Optimality and sub-optimality in a bacterial growth law. Nat Commun. 2017;8: 14123. doi:10.1038/ncomms14123
2. Li G-W, Burkhardt D, Gross C, Weissman JS. Quantifying Absolute Protein Synthesis Rates Reveals Principles Underlying Allocation of Cellular Resources. Cell. 2014;157: 624–635. doi:10.1016/j.cell.2014.02.033
3. Hui S, Silverman JM, Chen SS, Erickson DW, Basan M, Wang J, et al. Quantitative proteomic analysis reveals a simple strategy of global resource allocation in bacteria. Mol Syst Biol. 2015;11. doi:10.15252/msb.20145697
